# Designing Convergent Overlapping Genes with Transformer Encoder Models and Lightweight Structural Proxies

**DOI:** 10.1101/2025.11.07.687268

**Authors:** Jason K. Morgan

## Abstract

Overlapping genes allow multiple proteins to be encoded from a single DNA sequence, including convergent (antisense; tail-to-tail) orientations across three reading frames (phases 0, 1, and 2), with phase 1 most frequently observed in nature. Designing such overlaps is challenging due to codon degeneracy, phase-specific biases, and the need to preserve structural integrity for both proteins. Here, a purpose-built transformer encoder is introduced, trained on a balanced synthetic dataset of convergent overlaps spanning diverse prokaryotic genomes and GC contents. Controlled amino acid substitutions were incorporated during training to enhance model generalization, particularly for phase 1 overlaps. At inference, Monte Carlo dropout enabled uncertainty-aware sampling of synonymous codon solutions, which were iteratively refined using a windowed, multi-objective optimization framework. Candidate overlaps were scored using composite weighting across secondary structure preservation, substitution similarity, alignment identity, and ESM-2 contact map similarity, with the structural similarity index measure (SSIM) applied as a rapid proxy for structural fidelity. This approach generated convergent overlaps across all phases, with phase 1 showing the highest success rates. Optimization trajectories revealed distinct dynamics, with secondary structure preservation steadily increasing despite its lower weight. External validation using SwissProt proteins stratified by AlphaFold2 (AF2) predicted local distance difference test (pLDDT) confidence supported generalization to proteins with differing rigidity, yielding high secondary structure preservation in silico. These results demonstrate that transformer models trained directly at the nucleotide level, when coupled with uncertainty-aware inference and lightweight structural proxies, can support the computational design of synthetic overlapping genes without requiring full structural prediction. This framework offers a scalable path for phase-specific, codon-aware overlap design under realistic constraints.

## Introduction

The study of overlapping genes is an important area of interest in prokaryotic genetics, attributed to their ability to encode multiple functional products from a single DNA sequence (1–4). For example, a single gene sequence might code for two distinct proteins, each playing a different role in cellular function. Advancements in identifying and designing these overlapping sequences promise the creation of genetic constructs that are both more efficient and compact, potentially boosting the functionality and stability of engineered organisms (5,6). This phenomenon not only poses challenges but also opens new avenues in genetic engineering and synthetic biology (3).

Overlapping genes have been described using different approaches, but can be broadly grouped into unidirectional (sense; encoded on the same strand, → →) or convergent (antisense; encoded on the opposite strand) orientations (**Fig 1A**); convergent overlaps can be further refined into “head-to-head” (← →) or “tail-to-tail” (→ ←) orientations (7). Convergent overlapping genes occur in the same reading frame (phase 0), or frameshifted by one (phase 1) or two (phase 2) nucleotides. For example, in phase 2, convergent overlaps share their third (degenerate) codon positions. Research into convergent (tail-to-tail; antisense) overlapping gene sequences in prokaryotes has included analyses on the length distribution among overlapping and near-overlapping genes (8), investigations into their prevalence (and misannotations) within prokaryotic genomes (9,10), as well as the characterization of convergent gene pairs (11–18). Notably, both convergent and unidirectional overlaps exhibit a non-uniform distribution with a significant phase bias, predominantly favoring phase 1 overlaps (8,19,20). While the majority of convergent overlaps fall into phase 2, when excluding 4 nucleotide overlaps, nearly half are instead found in phase 1 (8). These studies collectively highlight the phase preference and distribution patterns of naturally occurring gene overlaps.

**Fig 1.**
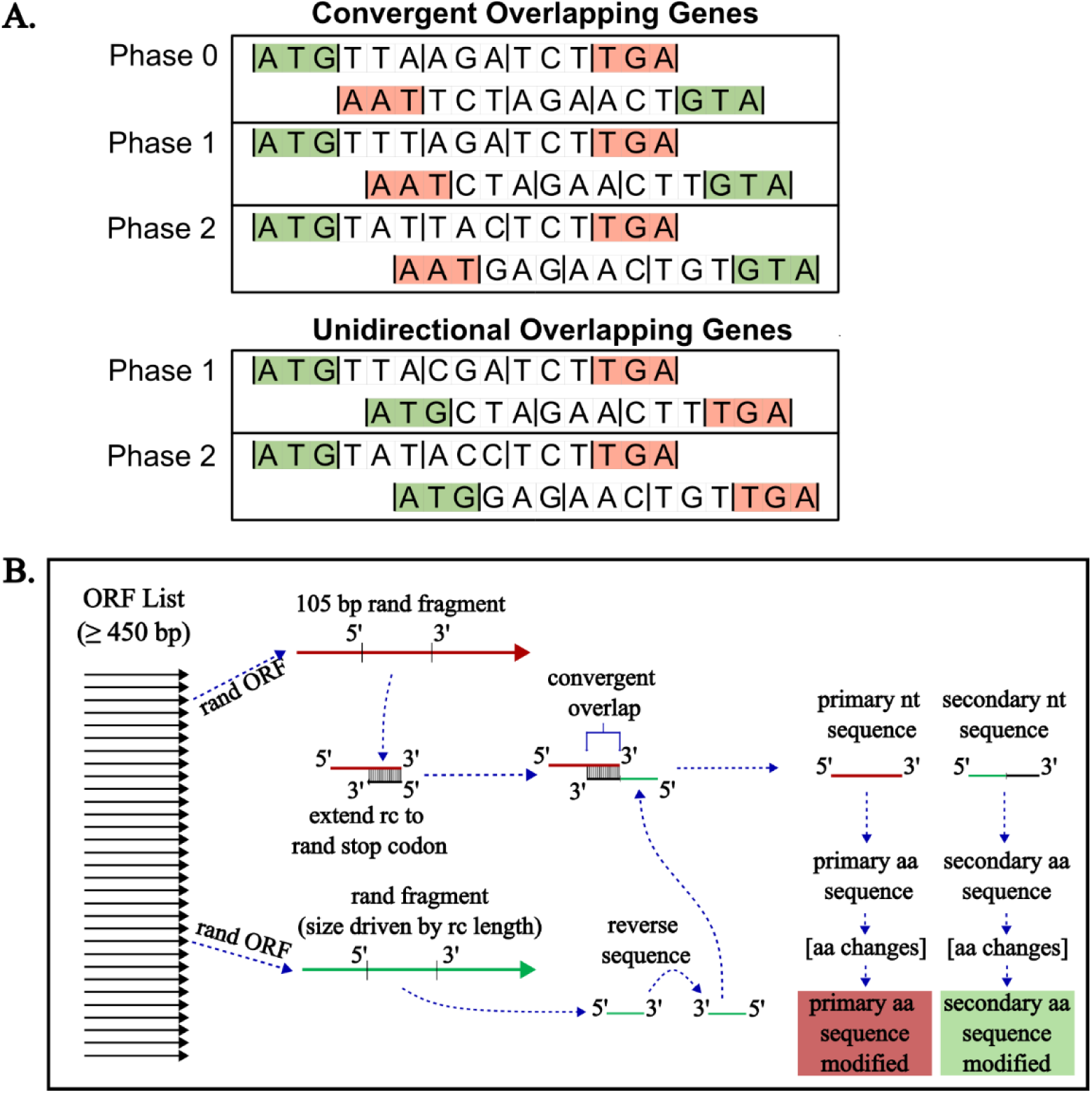
Depiction of convergent overlapping genes in three reading frames and the process developed to computationally design synthetic convergent overlaps from coding sequences. (A) Sequences are represented for convergent (tail-to-tail) and unidirectional (same-strand) overlapping genes in each of the three reading frames (phase 0, phase 1, and phase 2). Note, phase 0 unidirectional overlaps are not represented as these are considered as an alternative start site for the same gene (90). In each phase, the reading frame was shifted by one nucleotide. Start and stop codons are indicated with green and red shading, respectively. (B) The general process for creating synthetic convergent overlapping genes is presented, starting with a pool of ORF sequences filtered by length (≥450 base pair [bp]). The primary sequence and in-frame 105-bp fragment were randomly selected. A randomly selected stop codon from the reverse complement (rc), and the fragment reverse complement was extended to that stop codon to serve as the downstream sequence of the secondary sequence. Given the downstream secondary sequence length, another sequence was randomly selected from the same gene list. A size appropriate fragment was selected from that gene with length equal to the difference between the desired gene length and the upstream overlap fragment (ie, the reverse complement of the primary sequence to the selected stop codon). The upstream fragment was then combined with the downstream fragment to generate the complete secondary sequence. The resultant primary and secondary nucleotide sequences, sharing a known convergent overlapping region, were then translated to amino acid (aa) sequences (either modified or unmodified) for subsequent use in model training or inference.

The successful prediction of overlapping nucleotide sequences encoding two amino acid sequences poses unique computational challenges, and has been previously explored (6,21–25); the landscape of available methods has been summarized (22) with additional information pertaining to more recently released programs provided in **Supplementary Table 1**. An early solution for the design of unidirectional overlapping protein sequences included design of an algorithm to identify the shortest DNA sequence which could encode two amino acid sequences (26). More recently, examples included a dynamic programming algorithm (6) and a subsequent method that combines dynamic programming with a hidden Markov model (HMM) (5,27). The strategy was further extended to arbitrary pairs of natural protein domains (28). These approaches have collectively shown that it is possible to design fully embedded overlapping genes that encode functional proteins with high homology to starting sequences, underscoring the potential for engineered overlapping genes in prokaryotic genetics. Another approach, termed overlapping, alternate-frame insertion (OAFI) demonstrated the ability to design unidirectional overlaps through insertion of one gene into an alternate reading frame of another by targeting sites that may tolerate insertions (29). Recently, Byeon et al. (2025) demonstrated that deep generative models can be used to design synthetic overlapping genes spanning distinct protein families, achieving high in silico and experimental success rates even under the constraints of the standard genetic code (30). Their results indicate that sequence space may contain many viable overlapping solutions, suggesting broader applicability of computational approaches to both unidirectional and convergent overlaps.

Transformer models have demonstrated superior performance in various sequence modeling tasks due to their ability to capture long-range dependencies and contextual information (31–33). This study leverages a novel application of the transformer encoder-based model architecture to predict synthetic convergent overlapping genes in prokaryotes. Unlike dynamic programming approaches, transformers utilize self-attention mechanisms to process the entire input sequence simultaneously (32), enabling the model to capture patterns and dependencies that span across long sequences. In this study, Monte Carlo (MC) dropout is applied during inference to enhance prediction performance. Dropout, more generally, is a method of reducing overfitting in deep neural networks during training (34). MC dropout retains dropout activity during inference, effectively sampling from a distribution of possible model predictions (34–36). This provides a practical Bayesian approximation of model uncertainty (37), which is especially advantageous for predicting overlapping genes where the degeneracy of the genetic code permits many distinct but functionally viable nucleotide encodings. By sampling multiple plausible solutions, MC dropout enables exploration of the many synonymous coding possibilities inherent to the genetic code, increasing the likelihood of identifying overlaps that preserve the structural and functional integrity of both proteins.

The objectives of this study are to (i) evaluate the ability of transformer encoder models to recover convergent overlaps across all three phases, (ii) quantify the impact of overlap length and sequence variability on prediction success, and (iii) assess multi-objective optimization as an alternative to full structural prediction in improving overlap fidelity.

## Methods

### Synthetic Overlapping Gene Dataset Construction for Model Training

While naturally occurring convergent overlaps have been systematically assessed and documented (9), there is no readily available database of convergent overlapping genes with equally distributed overlap lengths. Given this limitation, a database of synthetic convergent overlaps was generated using a diverse set of open reading frames (ORFs) from prokaryotic genomes. Synthetic convergent overlapping genes were created using ORF sequences from 24 different genomes from prokaryotes (**Supplementary Table 2**), using the R (version 4.3.1) and Python programming languages (version 3.8.8). Genomes were selected to have a wide range of genomic GC content (ranging from 23.3% to 74.9%, with a mean of 52.41%), with the aim of controlling for potential amino acid compositional bias driven by GC content (38). The between-strain coding GC content (calculated using the set of ORF sequences from each strain) ranged from 20.7% to 72.7%, with a mean of 50.5%.

Briefly, for each strain, sequences were downloaded from the NCBI database (https://www.ncbi.nlm.nih.gov/genome/) and each FASTA file was processed to extract ORFs, which were stored in separate list for each strain (**Fig 1B**). ORF sequences were filtered to remove sequences shorter than 450 nucleotides, to exclude sequences with insufficient opportunity for varied random sampling during overlap construction. A function was defined to randomly select in-frame segments from the ORF sequence lists, to include a primary and secondary sequence either with or without a convergent overlapping region. First, a contiguous in-frame 312-nucleotide length section from each ORF sequence was extracted and used as the basis for the primary sequence. The primary sequence segments were modified to ensure they contained start and randomly selected stop codons, as these were removed during initial sequence processing.

The reverse complement of this synthetic gene was selected to generate the secondary sequence with a defined convergent overlap length. A uniform set of overlap lengths was generated for each length by extracting the reverse complement, adding a stop codon, and removing any in-frame stop codons with a random codon. Both the added stop codon and in-frame replacements were selected such that stop codons in the forward direction were not introduced. However, owing to the methodology employed, a portion of generated secondary sequences with overlapping sequences contained more than one stop codon and were subsequently filtered from the dataset to ensure all sequences contained only one stop codon and encoded a single amino acid sequence.

The amino acid sequence pairs with known overlaps were modified to incorporate up to 4 of BLOSUM62 weighted amino acid changes. The changes included altering amino acid sequences such that 1) the internal reverse stop codon was removed from both amino acid sequences (primary and secondary), while keeping the known overlap for the original amino acid sequences, or 2) the internal reverse stop codon was removed from both amino acid sequences (primary and secondary), while also introducing additional amino acid modifications per sequence. This yielded modified primary and secondary amino acid sequences, with an unmodified known convergent overlap nucleotide sequence capable of encoding amino acid sequences with high amino acid sequence identity to the primary and secondary amino acid sequences.

The length of the input amino acid sequences was limited to 104 amino acids to manage computational complexity during training and inference phases. Each amino acid was separated by a space and the overall included a terminal asterisk (*) and the primary and secondary sequences were concatenated to generate a single input sequence for training.

To balance the representation of different overlap lengths, data were stratified based on the length of the overlap. The overlapping gene training data were balanced, ensuring equal representation for each strain and each length of nucleotide overlap (specifically, convergent overlaps of length 199 to 312 nucleotides). The final dataset was composed of synthetic gene overlaps, each represented by a pair of modified amino acids sequences with a known target convergent overlap sequence. This dataset served as the training set for the transformer encoder-based models.

### Initial Model Description and Training

The synthetic overlapping gene dataset was used to train transformer-encoder based models with the aim of generating convergent (tail-to-tail) overlaps for any two specified amino acid sequences. A total of 8,892,000 combined pairs, corresponding to 78,000 pairs per convergent overlap length.

Transformers have demonstrated superior performance in various sequence modeling tasks, particularly where understanding long-range dependencies is important (31–33). As such, this study utilizes a PyTorch based transformer encoder model for overlap prediction (39). Tokenization and text vectorization were performed using the TensorFlow Keras API (40), and data were split into training and validation sets. During training, the models each utilized 156 embedding dimensions, 13 attention heads, 192 feed-forward network dimensions, 5 blocks, a dropout rate of 0.1, and a total of 802,983 trainable parameters (**Fig. 2D**). Models were compiled using the Adam optimizer with cross-entropy as the loss function.

**Fig 2.**
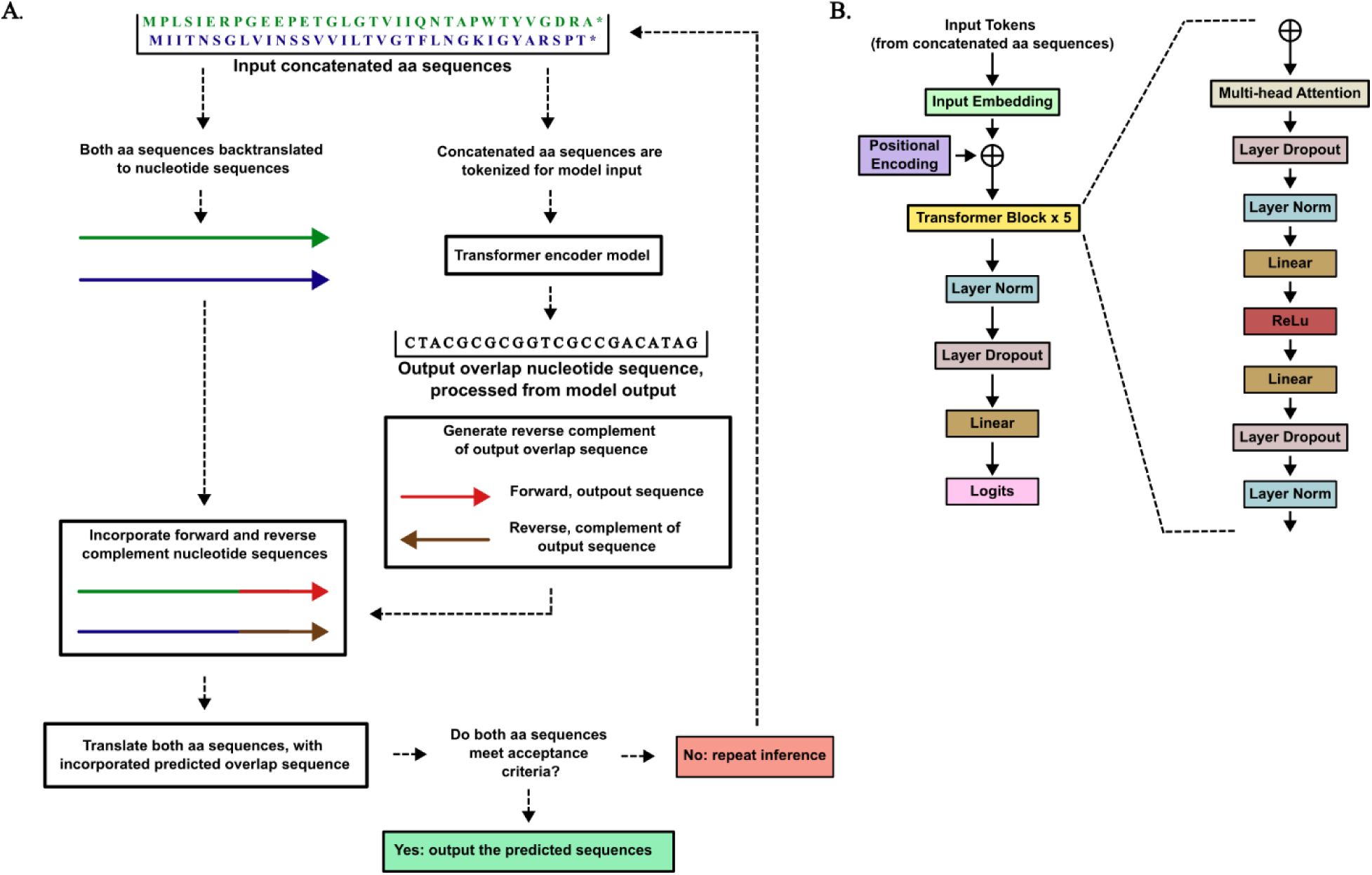
Illustration of the overlap transformer-encoder architecture and general structure of the data flow. (A) Depiction of the general structure of the data flow, for two input concatenated amino acid sequences (primary and secondary) used during inference. The primary and secondary amino acid sequences are represented with green and blue text, respectively, and the flow proceeds from top to bottom. (B) Illustration of the transformer encoder model architecture. The model input is tokenized concatenated paired amino acid sequences, and the model outputs raw logits. The output logits are processed using a greedy decoder to generate a predicted overlap sequence.

Each model was trained for 3 epochs with a batch size of 32 on a single NVIDIA GeForce GTX 5070 Ti GPU with 16GB of memory, and reached a validation accuracy of ≥90% by the third and final epoch.

### Test Dataset Generation and Model Inference

Amino acid pairs were generated as described above for preparation of the training dataset, but without the final modification step; however, a different set of strains (n = 12) were selected than those used for training, with a wide range of genomic GC content (ranging from 23.6% to 74.1%, with a mean of 50.2%) (**Supplementary Table 3**). Given that inference was performed using only coding sequences, the coding sequence GC content was calculated for each strain, and the corresponding mean GC value was used for subsequent analyses (varying from 24.2% to 72.1%, and a mean of 50.0%).

It is noted that inference was performed using the trained models along with the same tokenizer to vectorize the input sequence. Using a “greedy decoder” (41), the token with the highest probability at each step of the sequence generation process was selected.

MC dropout was implemented during inference to allow exploration of the uncertainty in model predictions (eg, variation in predicted overlap sequences), leveraging the benefits of Bayesian inference, as previously described (34,36,42). During inference, depending on the objective, feedforward dropout was set to 0.275 for initial predictions.

An algorithm was developed to generate overlapping gene sequences from input amino acid sequences using the trained models. The algorithm first back-translated the input amino acid sequences into a nucleotide sequence, leveraging *E. coli* codon usage frequencies to reflect the probabilistic distribution of codons for each amino acid (43). After back-translation, the models were used to predict potential overlaps within these nucleotide sequences. Predicted overlaps were then integrated into the original nucleotide sequences, resulting in a collection of modified sequences that incorporated the predicted overlapping regions downstream of the remaining back-translated nucleotide sequence. These modified sequences (ie, the back-translated and predicted regions) were then translated into amino acid sequences to assess the accuracy of the predictions. In instances where the correct amino acid sequence was not identified, additional rounds of prediction and back-translation were performed. This iterative process continued until either an overlap yielding amino acid sequences meeting the pre-specified criteria was identified, or the specified number of inference rounds were completed without a predicted overlap (defined as an unsuccessful prediction).

An alignment score was calculated for each sequence prediction. The predicted sequence was aligned against the known overlap using the Needleman-Wunsch algorithm (implemented via pairwise2 from Biopython (44)). For sequences with a known overlap and a predicted overlap sequence, the calculated alignment score was based on the length of the overlap, normalizing scores to a scale of 0 to 1 to facilitate comparison across predictions.

### Secondary Structure Prediction, Substitution Score, and Structure Prediction

The prediction of secondary structures was performed using S4PRED, which enabled predictions based on an input amino acid sequence alone without the need for multiple sequence alignments or known homologous sequences (45). S4PRED outputs were formatted as “ss2” files, from which the predicted secondary structure sequence was extracted for downstream analyses. The model employs a simplified three-state classification—α-helix (H), β-sheet (E), and coil/loop (C), which corresponds to a coarse-grained version of the DSSP (Dictionary of Secondary Structure of Proteins) annotations (46). Minor code modifications were implemented to enable efficient batch processing, substantially reducing runtime. These predicted structures were used to evaluate the structural compatibility of convergent overlapping gene candidates. Substitution matrices utilized were BLOSUM62 (47,48) and ProtSub (49). Protein structures were predicted either with AlphaFold3 or ColabFold (50–52).

### Optimization of predicted sequences

An iterative, windowed optimization procedure was used to convert paired amino-acid inputs into convergent (tail-to-tail) overlapping nucleotide sequences while preserving fold-relevant features of both proteins and maintaining sequence similarity. Input amino-acid sequences were first back-translated to an initial nucleotide scaffold using *E. coli* codon usage frequencies. The region targeted for overlap design was tiled into partially overlapping amino-acid windows; in the experiments reported here windows were 10 amino acids with an 8-amino-acid stride (10 AA window, 8 AA stride; model receptive field used = 105 AA). During optimization, only nucleotides mapping to the current window were permitted to change; flanking nucleotides were held fixed except at user-designated residues enforced by token-level biases (for example to preserve active-site residues or motifs). Bracketed constraints on the forward strand were mirrored on the reverse-complement strand so that enforced bases remained consistent with the convergent orientation.

Within each window a transformer-encoder was executed in an uncertainty-aware mode: Monte-Carlo (MC) dropout was retained at inference to generate diverse synonymous codon proposals. Token-level fixed-logit (base) biases were applied at specified nucleotide positions to enforce required bases during sampling; combined forward/reverse fixed-logit maps ensured constraints were respected on both strands. Each sampled nucleotide pair was translated to amino-acid sequences and duplicate amino-acid solutions were removed prior to scoring.

Structural compatibility was assessed using single-sequence secondary structure prediction (S4PRED; three-state H/E/C). Secondary structure preservation was reported as the percentage of residues for which a candidate’s predicted state matched the original sequence prediction. Substitution similarity was computed using BLOSUM62 and normalized to the original sequence’s self-score to yield a percent similarity; global alignment identity was also measured. ESM-2 (53) (model: esm2_t12_35M_UR50D) was used to generate residue-residue contact probability matrices, having demonstrated utility as a fast and effective single-sequence prediction feature (54). Contact map similarity was assessed using the structural similarity index (SSIM) (55), computed between ESM-2-predicted contact maps of candidate overlap sequences and their respective originals. SSIM quantifies local and global structural pattern agreement (range 0-1, with 1 indicating identical maps). Raw values were linearly scaled to a percentage (0-100) to yield the ESM-2 contact map score. This metric served as a rapid, computationally efficient proxy for structural preservation, enabling large-scale overlap design iterations while retaining sensitivity to perturbations in contact map topology.

Candidates were ranked by a fixed weighted composite score across all experiments: 0.15 (secondary structure preservation), 0.15 (substitution similarity), 0.1 (alignment identity), and 0.6 (ESM2 contact map similarity). Although this weighting scheme was not exhaustively optimized, empirical testing with randomly selected SwissProt proteins indicated that it yielded stable rankings across experiments and supported overlap generation while balancing structural fidelity with sequence similarity (**Figs. S3, S4, and S5**). The top-ranked candidate for the window was integrated, and optimization proceeded to the next window. Multiple full passes across windows were permitted for refinement. After the first structurally successful candidate, global feedforward and attention dropout levels were reduced to bias sampling from exploratory search toward fine-tuning, though the weighting scheme itself remained unchanged.

Efficiency measures implemented for reproducibility and speed included: (i) caching of secondary structure predictions so identical amino-acid sequences were scored only once, and (ii) caching of ESM-2 embeddings and contact maps. All candidate metadata (attempt, window, per-candidate metrics, and secondary structure predictions) were retained in memory and exported at the end of each input row to permit analysis.

### Mutual information, PCA, and robustness testing

Terminal regions (104 amino acids) from 250 randomly selected SwissProt-derived proteins were used as seeds to generate ensembles of mutated sequences. For each seed, trajectories were created by stepwise amino acid substitutions sampled from the 20-letter alphabet, introducing one mutation per step for up to 75 iterations. This yielded a progressive set of variant sequences per seed protein. Each sequence in the ensemble was then scored for global alignment identity, ESM-2 contact map similarity, and secondary structure match relative to the reference. These metric vectors formed the input for subsequent mutual information (MI) and principal component analyses.

MI was estimated after quantile discretization of continuous scores into 20 bins to evaluate pairwise dependence among objectives. To reduce sensitivity to binning, MI was also estimated using a k-nearest-neighbor approach. Statistical significance of discrete MI values was assessed using permutation testing with 1,000 random shuffles. Principal component analysis (PCA) was applied to z-scored metric vectors (alignment, ESM-2 contact map similarity, secondary structure match) to examine the dimensional structure of variation. Robustness checks included varying the number of bins (5–40) for MI and bootstrap resampling of PCA explained variance.

### Statistical Analyses and Figure Preparation

The R programming language (version 4.3.1) and the ggplot2 package (56,57) were used to perform statistical analyses and generate figures. Prediction success rates are reported as percentages with exact 95% confidence intervals calculated using the Clopper–Pearson method. Correlations between similarity metrics were assessed using Pearson’s correlation coefficient with associated *p*-values. Relationships between secondary structure composition (coil, helix, sheet) and prediction performance were examined using LOESS smoothing with 95–99% confidence bands. For external validation experiments, performance metrics were stratified by AlphaFold2 pLDDT confidence brackets and overlap phase. Group differences were evaluated using nonparametric pairwise contrasts with adjusted p-values (Bonferroni correction). Effect sizes are reported as absolute percentage-point differences between groups, alongside confidence intervals. Data are presented as mean values with ranges or confidence intervals where appropriate. Figures were generated with ggplot2, with additional overlays and schematics prepared in Inkscape (version 1.3.2).

### Declaration of Generative AI and AI-assisted technologies

During the preparation of this work, ChatGPT (5.0; OpenAI) was used to assess readability and language, and as a tool to generate code snippets and assist in debugging. After using this tool/service, the author reviewed and edited the content as needed and takes full responsibility for the content of the publication.

### Code and data availability

The datasets and code used for constructing the synthetic overlap dataset, training the models, and performing all analyses are available at: 10.5281/zenodo.17626094. Code for running model inference (to predict convergent overlaps) is available at: https://github.com/protosome/convergent_overlaps_aa_change

## Results

### Prediction of convergent overlapping genes from amino acid sequence pairs from a dedicated transformer-encoder model set

While prior work has established that it is technically feasible to generate overlapping genes from paired amino acid sequences, the performance of dedicated transformer-encoder models for this task has not been systematically evaluated. In this study, transformer-based models were trained to generate convergent overlapping nucleotide sequences from amino acid pairs, and their ability to recover overlaps across all three phases was assessed. These models were designed to identify convergent overlaps capable of encoding both input proteins, using recent advances in transformer-based sequence modeling to address the large solution space introduced by the degeneracy of the genetic code (31,58). Prediction quality was evaluated using alignment identity scores and secondary structure preservation metrics, allowing comparison across overlap lengths from 199 to 312 nucleotides.

Initial filtering was performed using a conservative alignment score cutoff of 34% (28,59). Without Monte Carlo dropout (ie, dropout enabled during inference), successful predictions were substantially reduced, and repeated inference runs converged to identical codon solutions. This indicated that deterministic decoding restricted the search space to narrow, low-diversity outcomes. By contrast, applying Monte Carlo dropout during inference expanded sequence diversity, producing multiple distinct codon assignments that encoded the same amino acid sequences. This stochastic sampling increased the likelihood of recovering overlaps that satisfied both similarity and structural thresholds, underscoring the importance of uncertainty-aware inference in navigating the combinatorial complexity of convergent overlap prediction.

To further assess training requirements, the effect of imperfect overlaps and amino acid substitutions in the training data was examined. Models trained only on unaltered overlaps exhibited markedly reduced success rates, particularly for phase 1 (**Figs. 3D, 3E**). In a shared test set of 5,000 amino acid sequence pairs without a known convergent overlap, training with modified sequences (stop codon removed plus 1–4 substitutions per sequence) substantially improved prediction rates: phase 0, 95.3% (94.7-95.9%); phase 2, 86.7% (85.7-87.6%); and phase 1, 81.6% (80.5-82.6%). In contrast, models trained on unaltered overlaps performed significantly worse: phase 0, 73.7% (72.4-74.9%); phase 2, 49.1% (47.7-50.5%); and phase 1, 9.7% (8.9-10.6%). These findings are consistent with the hypothesis that incorporating controlled sequence variation during training increased model flexibility and suggest improved generalization, particularly when coupled with Monte Carlo dropout during inference. On this basis, models trained with amino acid substitutions were selected for all subsequent analyses.

**Fig 3.**
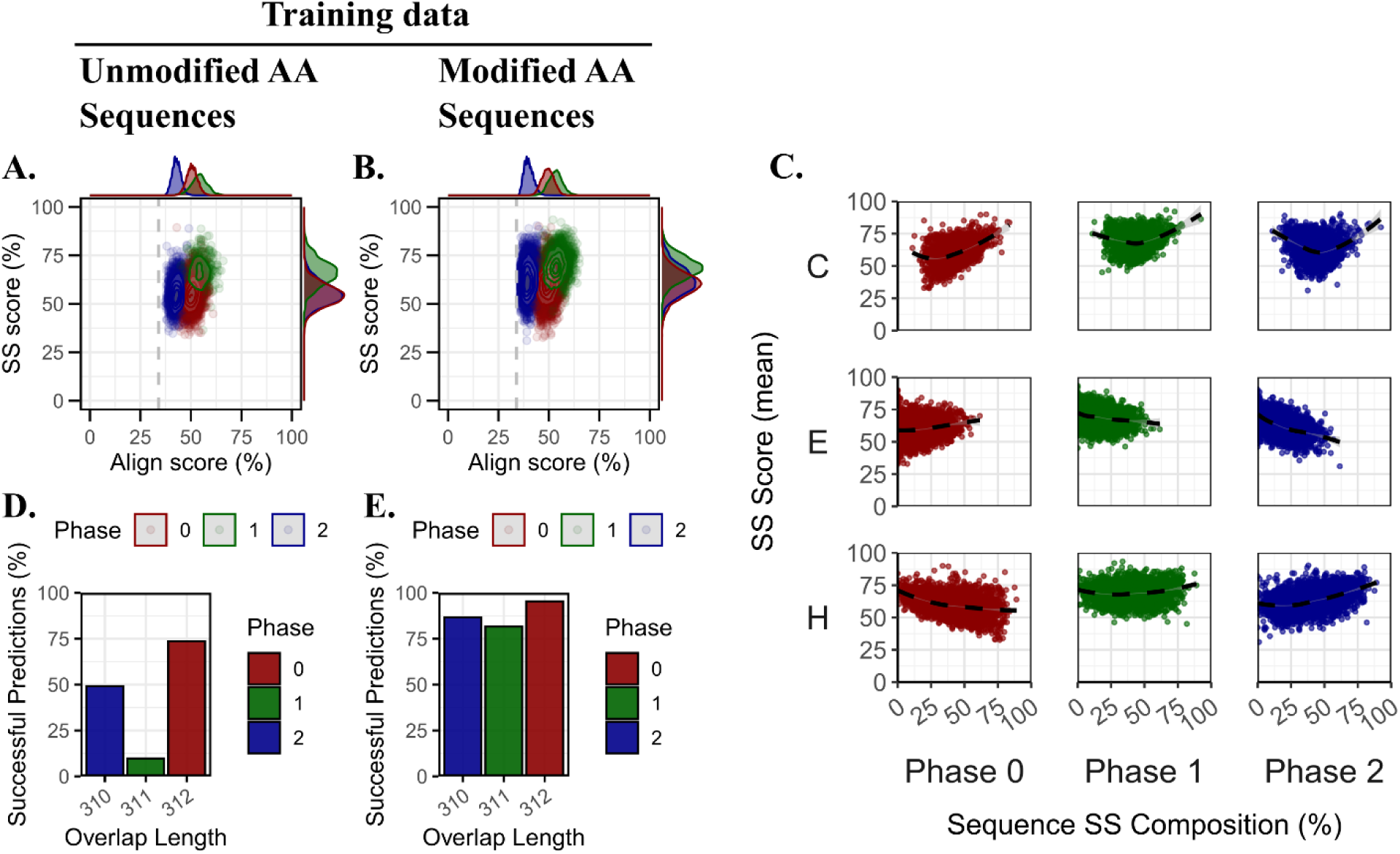
Training data evaluation of secondary structure preservation and convergent overlap-phase performance. (A, D) Results for unmodified amino acid sequences. (A) Scatter density plot of secondary structure (SS) score versus alignment score, with contours and marginal density plots showing the distribution by overlap phase (0 = red, 1 = green, 2 = blue). (D) Fraction of successful predictions across overlap lengths (310, 311, and 312 nucleotides), stratified by phase. Success rates are reported with exact 95% Clopper–Pearson confidence intervals. (B, E) Results for modified amino acid sequences. (B) Scatter density plot of SS score versus alignment score with the same phase-coloring scheme. (E) Fraction of successful predictions across overlap lengths for modified sequences, stratified by phase. Success rates are reported with exact 95% Clopper–Pearson confidence intervals. (C) Relationship between sequence-level SS composition (coil [C], sheet [E], helix [H]) and mean SS score across phases 0, 1, and 2. Dashed lines indicate LOESS regression fits with 99% confidence bands.

The outputs of these models were further evaluated using BLOSUM62 and ProtSub substitution matrices. For overlap lengths of 310-312 nucleotides, similarity scores were strongly correlated with amino acid identity (r = 0.88; p < 0.0001), and the matrices themselves were highly correlated with each other (r = 0.98; p < 0.0001).

In addition to substitution metrics, alignment scores and secondary structure scores were compared across phases. Phase 1 overlaps exhibited the highest average alignment and secondary structure scores, phase 0 displayed intermediate values, and phase 2 was generally the lowest. However, score distributions overlapped considerably, suggesting that phase-dependent differences were not absolute. The near-identical secondary structure score distributions for phase 0 and phase 2 further indicated that structural outcomes may be more sensitive to sequence-specific context than to phase alone.

To broaden the analysis, overlap lengths from 199 to 312 nucleotides were examined across a larger set of amino acid pairs. Phase-dependent patterns persisted: phase 1 overlaps were increasingly recoverable at shorter lengths, and the relative rank order of secondary structure and alignment scores across phases remained consistent. These findings suggest that phase 1 overlaps may be predicted to be structurally and codon-compatible favorable, and that performance of the transformer encoder model set is influenced by overlap length and the diversity of sequences available for training.

### High Secondary Structure Scores Are Linked to Coil-Dominated Precursor Sequences

To assess whether the secondary structure composition of precursor amino acid sequences influences the predicted quality of convergent overlaps, the proportion of coil (C), β-sheet (E), and α-helix (H) structures in each input was compared against the model’s predicted secondary structure score. For each amino acid pair, structural proportions were calculated separately and then averaged across both sequences. This approach reflects the biological premise that overlap feasibility arises not from a single sequence in isolation, but from the combined structural constraints and flexibilities of the two proteins that must be co-encoded (60,61).

Across all three overlap phases, sequences with higher coil content exhibited elevated secondary structure scores (**Fig. 3C**). This trend was most apparent in phase 0 and phase 1, where LOESS-smoothed curves showed a modest increase in secondary structure score with increasing coil content, followed by a sharper rise beyond approximately 70% coil. In phase 2, a similar increase was observed at high coil levels, but the trend was less consistent, with greater variability at intermediate values. By contrast, α-helix content was inversely associated with secondary structure score. The most pronounced effect occurred in phase 0, where increasing helix content corresponded to a marked decline in secondary structure scores. A similar, though less steep, trend was observed in the other phases. For β-sheet content, no consistent relationship with secondary structure score was observed; across all three phases, smoothed curves were largely flat, indicating that sheet content neither strongly promoted nor hindered predicted structural quality.

To evaluate whether these relationships extended to sequence-level similarity, mean alignment scores were also plotted against precursor structural compositions. No meaningful trends were observed for any secondary structure class (**Fig. S1**). Alignment scores remained relatively constant regardless of coil, β-sheet, or α-helix content, suggesting that precursor structure did not systematically influence the model’s ability to recover the reference overlap sequence.

### Multi-objective optimization and computational design of EGFP/AmpR convergent overlaps

Although the transformer-encoder models were capable of generating convergent overlaps for paired amino acid sequences, the secondary structure preservation achieved in initial predictions was variable and generally lower than values reported in prior work (60). To address this limitation, an iterative sequence optimization algorithm was implemented to refine model outputs using a multi-objective framework that did not rely on MSA availability (**Fig. 4A**).

**Fig 4.**
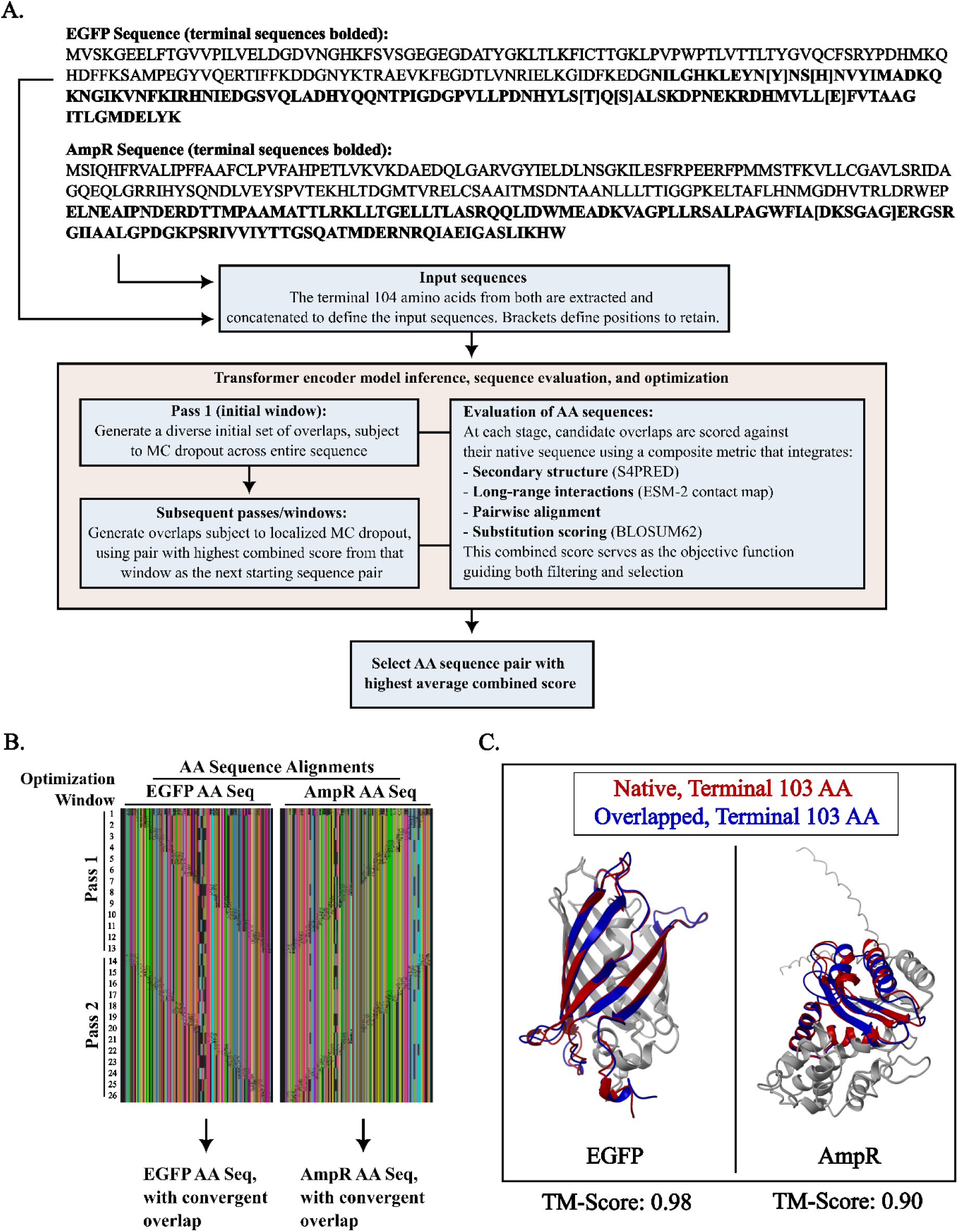
In silico design of convergent overlap with EGFP and AmpR. Overview of the multi-objective optimization process for EGFP and AmpR proteins using the transformer encoder framework. (A) Schematic of the transformer encoder–based workflow. The terminal 104 amino acids of EGFP and AmpR (across all tested overlap lengths) are concatenated and passed through the model. Outputs are scored by a composite metric integrating alignment, substitution, secondary structure preservation, and long-range interactions. Bracketed amino acids from the input are preserved to enforce positional constraints. The final sequence pair is selected from the output table by the highest average combined score. (B) Multiple-sequence alignment of all output pairs (n=3,624) for EGFP and AmpR across both optimization passes and windows (top to bottom). Diagonal dropout patterns correspond to localized MC dropout applied to the overlap region. Variability reflects bracket-constrained codon degeneracy introduced on the non-fixed partner sequence. (C) Predicted tertiary structures generated with AlphaFold3 for EGFP (left) and AmpR (right). Native and overlapping terminal regions relevant to the 311-nt overlap are highlighted (native in red, overlapped in blue); non-overlapping regions are shown in gray.

The optimization procedure employed a windowed feedforward Monte Carlo (MC) dropout strategy to generate multiple candidate solutions for each sequence window (**Fig. 4B**). Candidate overlaps were then evaluated against two complementary structural criteria: 1) predicted secondary structure states derived from S4PRED (62), providing local fold-relevant constraints, and 2) ESM-2 contact map similarity (63), which encodes sequence-level context that may reflect long-range residue dependencies. A key feature of the framework was the ability to enforce selective preservation of amino acid residues when required. This was achieved by applying a feedforward mask to specific codon positions, constraining the model to maintain defined residues (for example, active-site motifs) while permitting synonymous variation elsewhere. In practice, this allowed critical sequence features to be retained while enabling exploration of codon-level degeneracy to optimize overlap formation.

Relationships among the chosen objectives were next quantified using mutual information (MI) and principal component analysis (PCA). Both approaches indicated that alignment identity, ESM-2 contact map similarity, and predicted secondary structure capture overlapping but distinct constraints. Discrete MI (20 quantile bins) showed that alignment carried moderate information about both ESM-2 contact map similarity (MI = 0.393) and secondary structure match (MI = 0.381), whereas the MI between ESM-2 contact map similarity and secondary structure similarity was lower (MI = 0.240), consistent with partial independence. Continuous MI estimates (kNN regression) produced similar results (0.509, 0.508, and 0.340, respectively). All associations were highly significant in permutation tests (p = 0.001). PCA of z-scored metrics supported this interpretation (**Fig. S2**). A single axis (PC1) explained 72.4% of variance (95% CI: 71.8-73.0) and loaded positively on all three objectives, reflecting a general similarity dimension. PC2 (18.4%; 95% CI: 18.0-18.9) contrasted ESM-2 contact map similarity with secondary structure similarity, while PC3 (9.2%) primarily captured sequence-only variance. Thus, although alignment dominates the shared signal, secondary structure preservation and ESM-2 contact map-based similarity provide orthogonal information that cannot be reduced to alignment alone. This justifies their inclusion as separate objectives during overlap optimization.

Metric behavior under progressive sequence divergence was also assessed by subjecting 250 randomly sampled SwissProt proteins to iterative single-residue substitution trajectories (75 steps per sequence). Across trajectories, all four similarity measures declined with increasing mutational distance, but the rate and pattern of decay differed by metric (**Fig. S3**). Alignment identity and BLOSUM62 similarity decreased rapidly and near-linearly, consistent with their direct dependence on residue-level identity. By contrast, ESM-2 contact map similarity and secondary structure similarity showed more gradual declines. These distinct decay profiles support the interpretation that the objectives capture complementary aspects of overlap fidelity.

Weighting experiments further clarified the contributions of each objective. When the relative weights assigned to secondary structure similarity, ESM-2 contact map similarity, alignment identity, and BLOSUM62 substitution similarity were varied across randomly selected SwissProt pairs with very high AlphaFold2 pLDDT values (≥90), distinct trade-off dynamics were observed (**Fig. S4**). Increased weighting on secondary structure similarity consistently improved secondary structure preservation but reduced alignment and substitution similarity, whereas embedding-focused weightings enhanced ESM-2 contact map similarity at the expense of secondary structure similarity. Intermediate combinations (eg, 0.15-0.4 secondary structure with 0.6 ESM-2 contact map similarity) yielded more balanced outcomes across the four metrics. Variance patterns indicated that secondary structure weighting stabilized optimization trajectories, while ESM-2 contact map similarity remained comparatively variable across candidate sequences.

A convergent overlap between enhanced green fluorescence protein (EGFP) and a TEM-1 β-lactamase (ampicillin resistance marker from pCVD004; hereafter AmpR) was designed and refined under the same objectives as a biologically relevant test case (64,65). The combination of MC dropout sampling, secondary structure prediction, and embedding-based scoring consistently improved fold preservation of computationally designed EGFP/AmpR 311 nucleotide (phase 1) overlaps (**Fig. 4C**; **Fig. 5**). Compared to unoptimized outputs (ie, sequences generated from the first window), optimized sequences displayed higher secondary structure similarity rates and increased ESM-2 contact map similarity, demonstrating that overlap designs can be improved using lightweight, MSA-independent objectives. During windowed optimization, alignment and BLOSUM62 metrics remained largely constant or decreased across windows, reflecting their role in constraining candidate sequences to a biologically plausible subset of sequence space. In contrast, secondary structure similarity generally increased throughout optimization, despite being weighted at only 0.15 in the composite score. ESM-2 contact map similarity increased modestly in the early windows before plateauing. These findings suggest that fold-relevant secondary structure metrics may act as the principal discriminative driver of optimization progress, with alignment- and embedding-based metrics providing a stabilizing baseline. Following optimization, a pair of amino acids with the highest combined score was selected for structural comparison using predictions from AlphaFold3 (50), while selecting amino acid sequences in both pairs for preservation based on predicted importance (eg, active site). The AlphaFold3-predicted models of designed overlaps were highly similar to reference predictions (**Fig. 4C**), with TM-scores of 0.98 (EGFP) and 0.90 (AmpR), the latter with deviations generally localized to a disordered coil not located in the overlapping terminal region (66).

**Fig 5.**
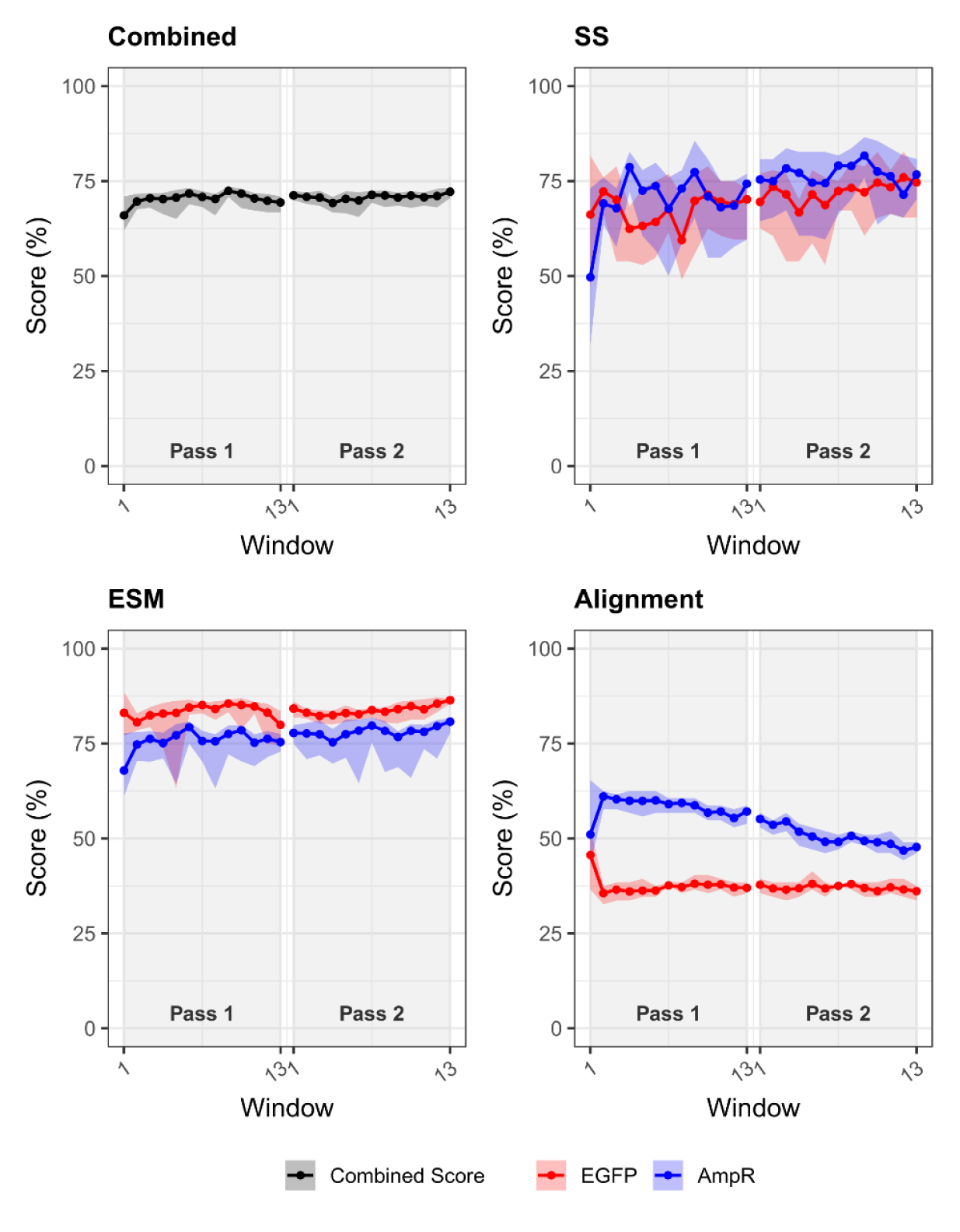
Trajectories of composite and component metrics during computational optimization of EGFP/AmpR convergent overlaps. Optimization performance for EGFP and AmpR across two passes, each consisting of 13 windows. Metrics shown include the combined score (top left), which integrates secondary structure (SS), ESM-2 contact map similarity, and alignment components, as well as the individual SS (top right), ESM-2 (bottom left), and amino acid alignment (bottom right) scores. Solid red (EGFP) and blue (AmpR) lines denote mean values, and shaded regions represent the minimum-maximum range across sequences within each window.

### Performance of this algorithm in predicting overlapping genes using pLDDT as an orthogonal proxy for intrinsic structure rigidity

Analyses on prokaryote-derived amino acid sequence pairs indicated that secondary structure preservation increased with coil content and decreased with helix content, with sheet content showing no consistent effect. To determine whether this pattern reflected a generalizable property of overlap design for convergent overlap derived from terminal amino acid sequences, an external validation set was assembled from SwissProt proteins stratified by AlphaFold2 (AF2) per-residue confidence (pLDDT), an orthogonal proxy for intrinsic structural rigidity (52,67–69). SwissProt sequences were filtered to 105–200 amino acids to align with the 105-AA receptive field and to bound computational cost, then binned into Low-intermediate (50–69), Confident (70–89), and Very high (90-100) pLDDT groups. For each bracket, 100 non-redundant amino-acid pairs were formed, and convergent overlaps were designed across phases 0, 1, and 2 at 310, 311, and 312 nt using the same transformer-encoder, windowed Monte-Carlo (MC) dropout inference, objective weights, and constraint handling described previously.

Quantitative analysis of secondary structure composition (determined using S4PRED) across pLDDT confidence bins revealed clear trends. Looking specifically at the terminal AA regions, as pLDDT confidence increased from the Low-intermediate to the Very high bins, coil content declined substantially (from ∼64% to ∼41%), while both helix and sheet content rose (helix from ∼29% to ∼39%; sheet from ∼6% to ∼20%) (**Figs. 6A, 6B**). These shifts suggest that AlphaFold’s higher-confidence regions correspond generally to more ordered structural states, and reinforce the interpretation of pLDDT as an approximate proxy for structural order and rigidity (50,70,71).

**Fig 6.**
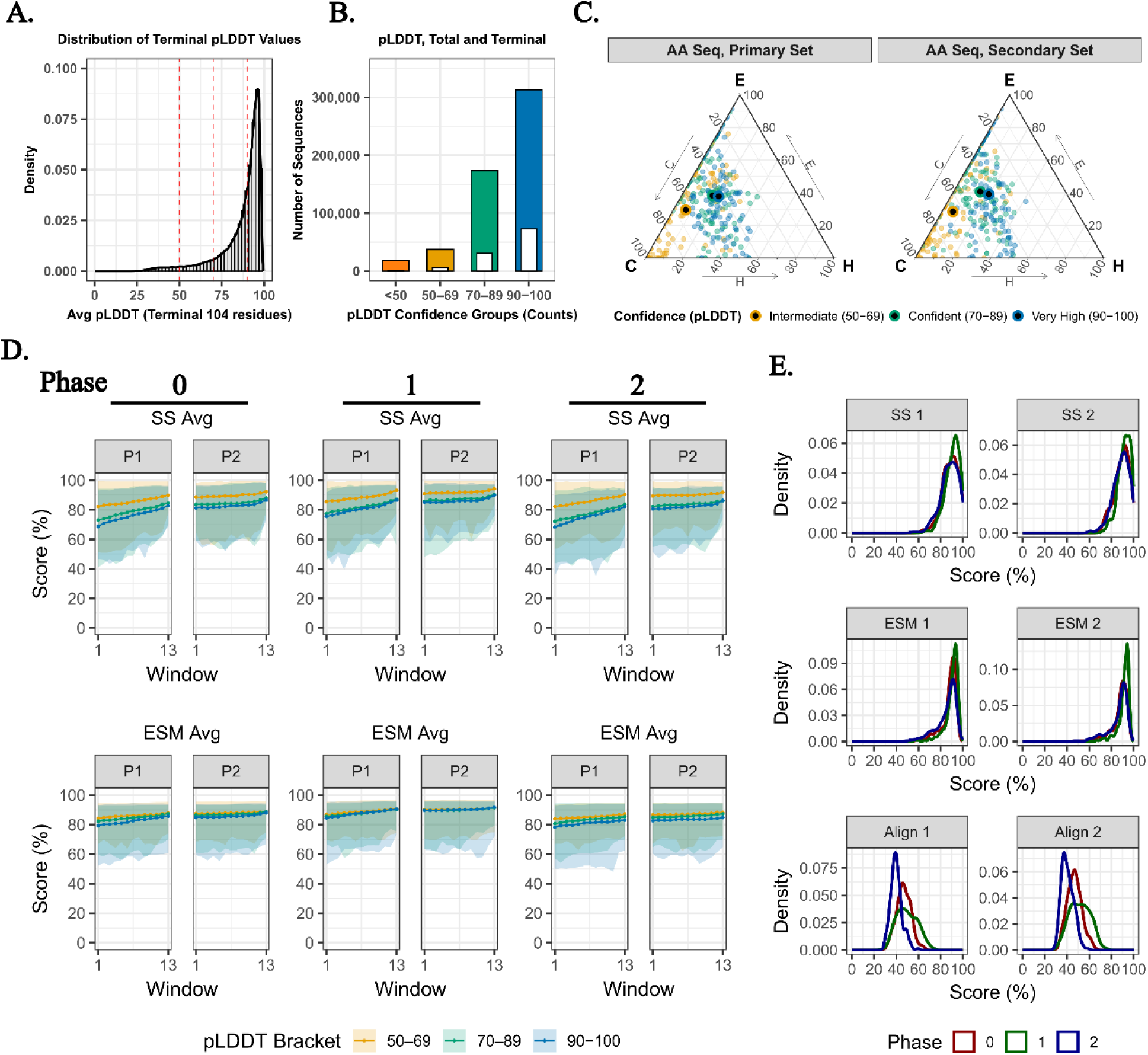
Comparison of computational overlap predictions across pLDDT brackets, with structural composition and optimization outcomes. Analyses are based on random sampling of SwissProt protein pairs (n=100 for each bracket) from the AlphaFold2 Protein Structure Database (AF2). (A) Distribution of average pLDDT values for the terminal 104 residues across sampled sequences, with red dashed lines marking the confidence thresholds. (B) Counts of sequences assigned to pLDDT confidence brackets, shown for both total and terminal regions (internal white bars denote terminal regions). (C) Ternary plots of amino acid composition (helix [H], sheet [E], and coil [C]) for the primary and secondary sequence sets, colored by pLDDT bracket (low-intermediate, confident, and very high). Circles with black centers denote the centroid (mean position of all sequences within a given pLDDT bracket) in ternary space. (D) Optimization performance across convergent overlap phases (0, 1, 2) for Pass 1 (P1) and Pass 2 (P2). Plots show average secondary structure (SS Avg) and ESM-2 contact map similarity (ESM Avg) scores by optimization window, with shaded regions representing the range of values stratified by pLDDT bracket. (E) Density distributions of individual metric scores (SS, ESM, and Alignment) for the highest combined score sequences (primary and secondary) across phases 0, 1, and 2.

These AF2-stratified findings were consistent with the earlier composition-based analysis using prokaryotic ORFs in this study. Sequence pairs with lower pLDDT (coil-enriched) were associated with slightly higher secondary structure preservation, whereas higher-confidence inputs tended to show reduced variation (**Fig. 6C**). Nevertheless, the overall high secondary structure scores across all three brackets emphasize that the algorithm was able to recover structurally faithful overlaps even for proteins predicted by AlphaFold2 to adopt stable, well-defined folds. Alignment identity and substitution similarity remained generally stable across brackets, indicating that secondary structure preservation was the primary dimension along which flexibility exerted its effect.

Bracket- and phase-specific effects were evident across all three metrics, consistent with the secondary structure composition of the pLDDT bins. Across overlap phases, sequences in the Low-intermediate (50–69) bracket generally outperformed those in the Confident (70–89) and Very high (90-100) brackets, with the gap being most pronounced in phase 2 (**Fig. 6D**). In phase 1, the bracket effect was clear for secondary structure preservation and for the combined score relative to Very high (90-100), whereas pairwise differences were not evident for ESM-2 contact map similarity; Low-intermediate (50–69) also did not differ from Confident (70–89) for the combined score. Overall, phase 1 maintained the highest preservation, while phase 2 exhibited the largest bracket separations. Although group differences were statistically significant, effect sizes varied by metric: within a phase, combined and ESM-2 contact map similarity differences were typically ∼1-4 percentage points, whereas secondary structure differences reached ∼5-8 points in some phase 2 contrasts. All groups maintained mean secondary structure preservation above 85%. Full pairwise estimates with confidence intervals and adjusted p-values are provided in **Supplementary Tables 4 and 5.**

While statistically significant differences were observed between pLDDT brackets, these differences were relatively small, and overall secondary structure preservation remained consistently high across all conditions (**Fig. 6E**). These results reinforce the generalizability of the earlier coil-associated trend, but also highlight the robustness of the multi-objective, uncertainty-aware optimization framework in achieving high predicted structural preservation regardless of precursor rigidity.

## Discussion

Overlapping gene architectures have long been recognized as both a source of evolutionary constraint and an avenue for novel functionality (72–80). In addition to their evolutionary importance, overlapping structures can be purposefully engineered under realistic codon and structural constraints. The present study shows that phase-specific convergent overlaps can be computationally generated and optimized using a transformer encoder-based multi-objective framework without requiring full 3D prediction at the optimization stage; targeted 3D checks were applied post hoc. This framework offers a complementary tool that could expand the range of strategies available for synthetic overlap design, particularly where phase, length, and nucleotide-level control are critical.

Recent work has demonstrated that deep generative protein models can discover viable overlapping solutions under genetic code constraints. Byeon et al. (2025) showed that pretrained amino acid-space generative models can design synthetic overlapping genes across multiple reading frames, validating expression experimentally using Gibbs sampling with codon-compatibility constraints (30). Xu et al. (2025) further demonstrated that pretrained generative protein language models (ESM-3) can yield viable entangled protein pairs through structure-conditioned inverse folding and CAMEOS-based entanglements, filtered by cross-entropy and Potts energy scores (81). Importantly, Xu et al. established that pretrained generative models are capable of identifying functional overlap-compatible solutions, albeit in a limited experimental context (the InfA/AroB system) and without explicit codon-level optimization.

The framework presented here builds on this momentum but takes a different route. Rather than relying on pretrained amino acid-space sampling, it employs a purpose-built, phase-generalizable transformer encoder model set trained on a balanced synthetic dataset of convergent overlaps spanning diverse GC contexts and overlap lengths. Multi-objective optimization is integrated directly into the inference loop, with candidate overlaps iteratively refined through a windowed, uncertainty-aware (MC dropout) search. Evaluation criteria span predicted secondary structure preservation (S4PRED), amino acid substitution similarity (BLOSUM62), pairwise alignment identity, and ESM-2 contact map similarity. This approach emphasizes codon-level control while maintaining structural fidelity, enabling overlap recovery without requiring full 3D structural prediction at each iteration.

MC dropout was retained during inference to enable stochastic sampling of synonymous codon solutions, thereby broadening the effective search space (35). Candidate overlaps were ranked using a fixed composite score that balanced structural preservation, substitution similarity, alignment identity, and embedding-based metrics. This weighting scheme served as a practical approach for navigating competing design constraints without requiring full structural prediction at each iteration, and was most closely aligned with prior work arguing that synthetic overlap design requires explicit balancing of competing biological constraints and objectives (5,60). While other multi-objective optimization strategies could be considered in future work (such as ε-constraint methods (82,83), adaptive scalarization (84,85), or evolutionary multi-objective optimizers (86)), the fixed weighted framework proved sufficient to achieve high overlap fidelity under the conditions evaluated.

The synthetic convergent overlapping gene dataset construction method developed as part of this study provides a structured framework for exploring overlapping gene design in silico, as it permits generation of diverse and uniformly distributed overlaps. However, one potential limitation of this approach is the reliance on synthetic, rather than natural, overlapping genes for model training, which were generated using ORFs from a diverse set of prokaryotic strains. This may introduce a bias affecting the predictive ability and variability in sequence output. Despite this concern, the method employed to generate synthetic overlapping genes is in principle analogous to the formation of new overlaps through the process of gene extension following the loss of a stop codon (8). This results in synthetic overlapping genes where the second (extended) strand’s encoded amino acid composition is derived from the reverse complement of the first strand, without undergoing further selection. As recently reported, there is extensive evidence suggesting functionality in prokaryotic convergent (antisense) proteins, indicating that non-coding RNA regions could also encode functional proteins (87). For same-strand overlaps, a significant difference in overall composition compared to non-overlapping genes has been previously observed (88), along with a bias towards disorder-promoting amino acids (89). The extent to which this affects convergent overlapping genes remains unclear. Future work will therefore aim to (i) characterize compositional differences between natural and synthetic overlaps, (ii) extend design to longer proteins beyond the 199-312 nucleotide range considered here, and (iii) experimentally validate whether in silico-designed overlaps from this framework maintain expression, folding, and function in vivo.

In summary, this study establishes that phase-aware transformer models trained directly at the nucleotide level, when coupled with uncertainty-aware inference and multi-objective optimization, can recover computationally designed convergent overlapping genes under realistic genomic constraints. By integrating codon-level sampling with structural and substitution-based objectives, this framework complements recent amino acid-space generative approaches while offering length- and phase-specific control. These findings provide a computational foundation for experimentally testable designs and point toward a scalable strategy for compact, synthetic gene architectures.

