## Supplementary Figures for "Designing Convergent Overlapping Genes with Transformer Encoder Models and Lightweight Structural Proxies"

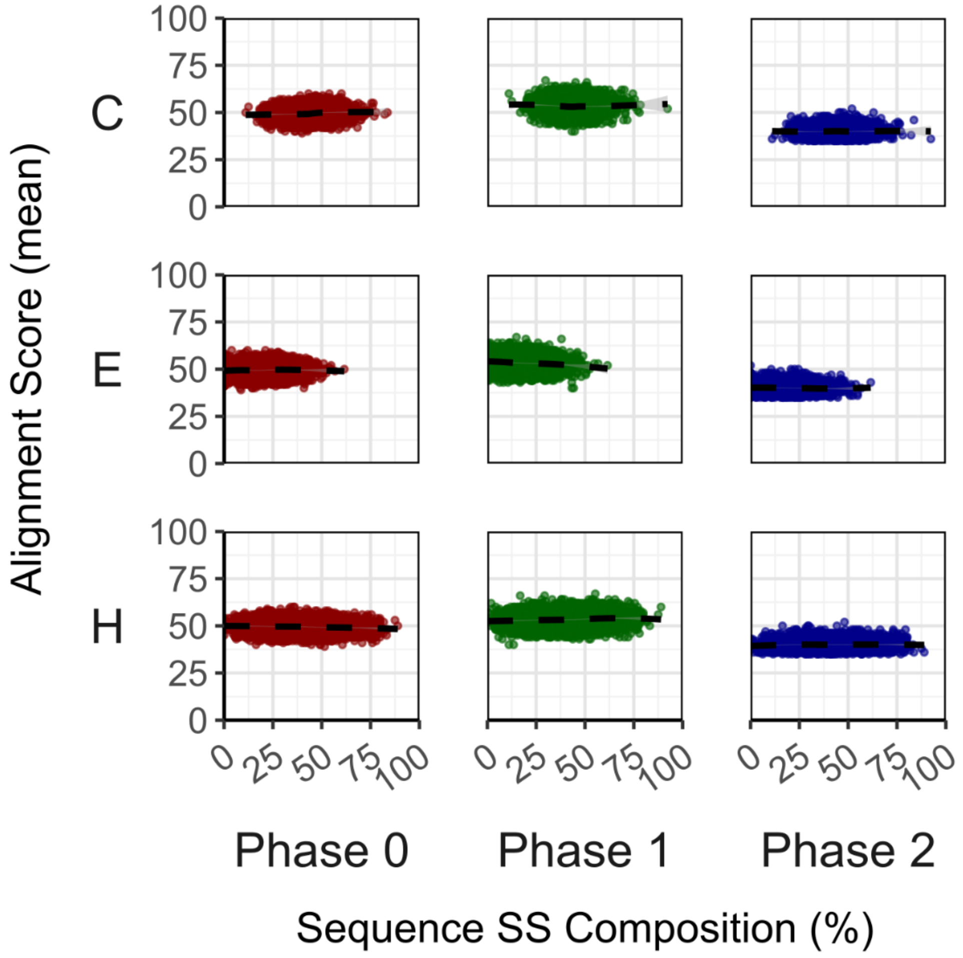


**Fig S1. Evaluation of data alignment performance for unoptimized convergent overlaps.**

Scatter plots showing alignment score (mean of both sequences) versus secondary structure (mean of both sequences; coil, strand, helix) composition across convergent overlap phases 0, 1, and 2. Alignment scores remain stable across structural compositions and phases.

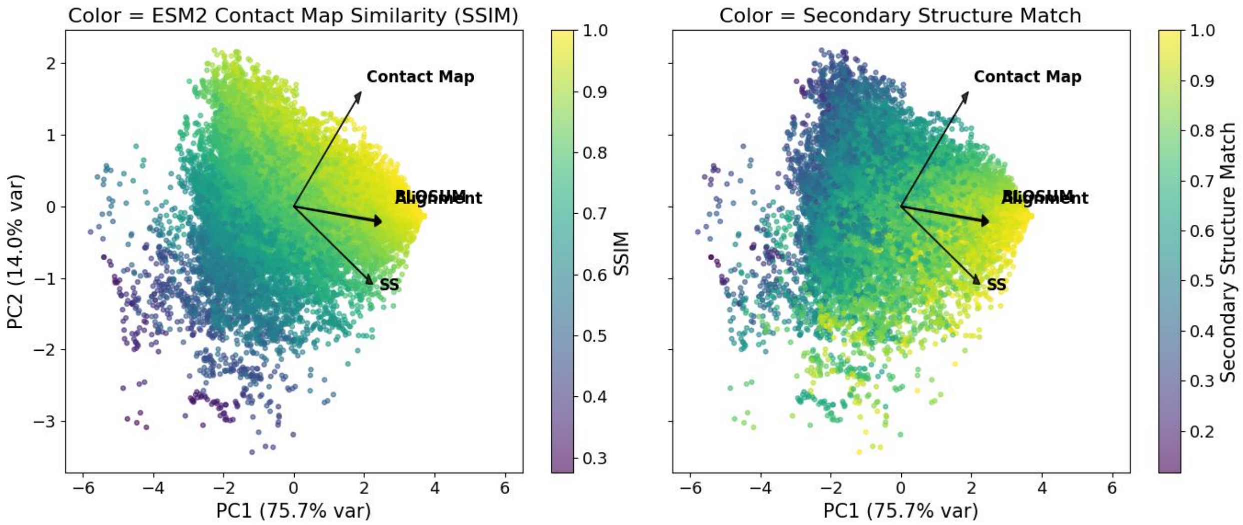


**Fig S2. Principal component analysis of similarity metrics for convergent overlap design.**

Principal component analysis (PCA) of alignment identity, BLOSUM-normalized substitution similarity, ESM2 contact-map similarity, and secondary-structure match. Each point represents an individual sequence, colored by secondary-structure match. Arrows indicate variable loadings on PC1 and PC2, highlighting shared and orthogonal contributions across the four objectives.

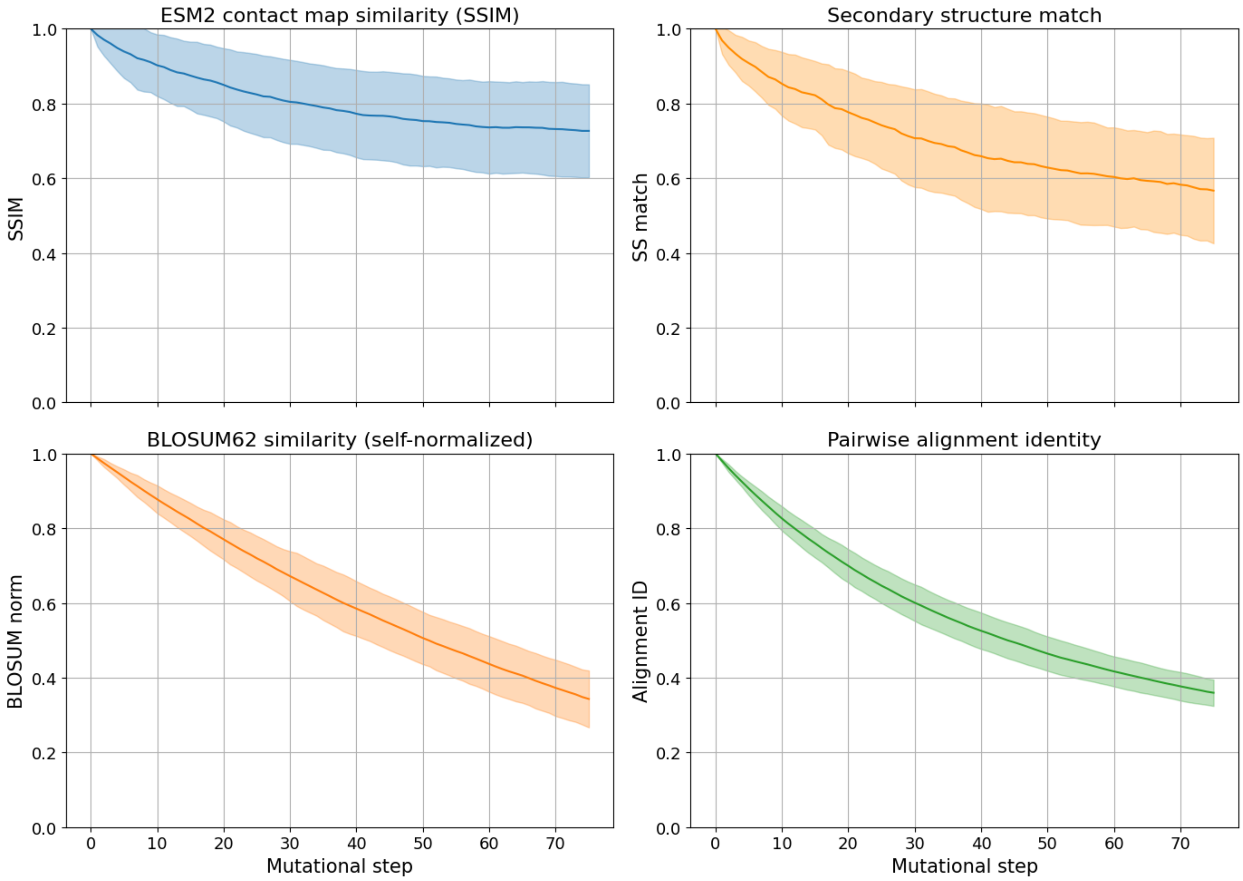


**Fig S3. Trajectory-level metrics for randomly sampled SwissProt proteins under iterative mutation.**

250 proteins were randomly sampled from SwissProt database, without replacement (N = 250). For each sequence, the terminal 104 amino acids were extracted, and a trajectory of 75 steps was generated by introducing one random amino acid substitution per step. At each step, similarity to the unmutated reference was quantified using four metrics: (top left) ESM2 contact-map similarity (SSIM), (top right) S4PRED-predicted secondary-structure match, (bottom left) normalized BLOSUM62 similarity, and (bottom right) pairwise alignment identity. Curves show mean ± standard deviation (SD) across all sampled trajectories.

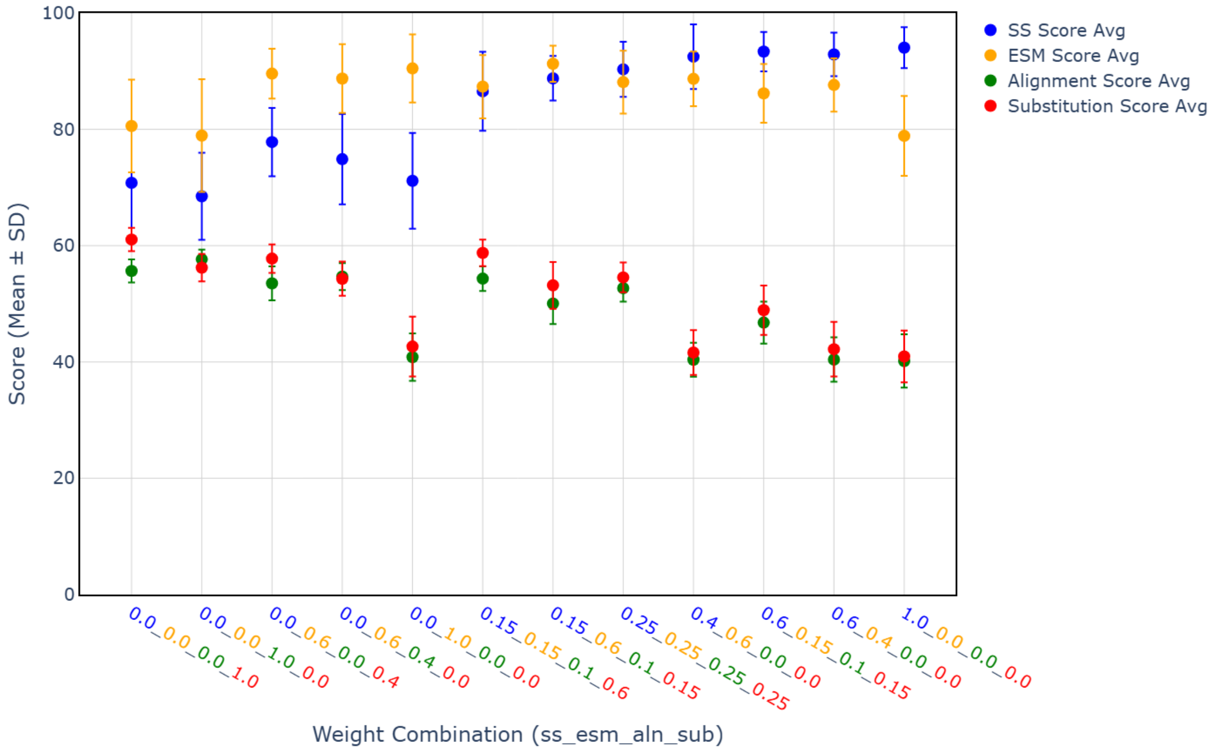


**Fig S4. Convergent overlap optimization metrics across parameter weightings using 15 randomly selected protein pairs from SwissProt.**

For each parameter weighting, the sequence pair with the highest combined score was identified, and the mean metric value across 15 optimization runs (n=15) was plotted. Error bars indicate ± standard deviation (SD). Four metrics are shown: secondary structure similarity, ESM-2 embedding similarity, alignment identity, and substitution similarity. Parameter combinations are denoted as ss_esm_aln_sub (eg, 0.15_0.6_0.1_0.15), corresponding to the relative weights applied to each scoring component. Only sequences whose terminal region exhibited a mean pLDDT ≥ 90 in the original AlphaFold2 prediction were included. This cutoff was applied to restrict the analysis to terminal regions predicted with very high confidence, thereby reducing potential noise from low-confidence residues.

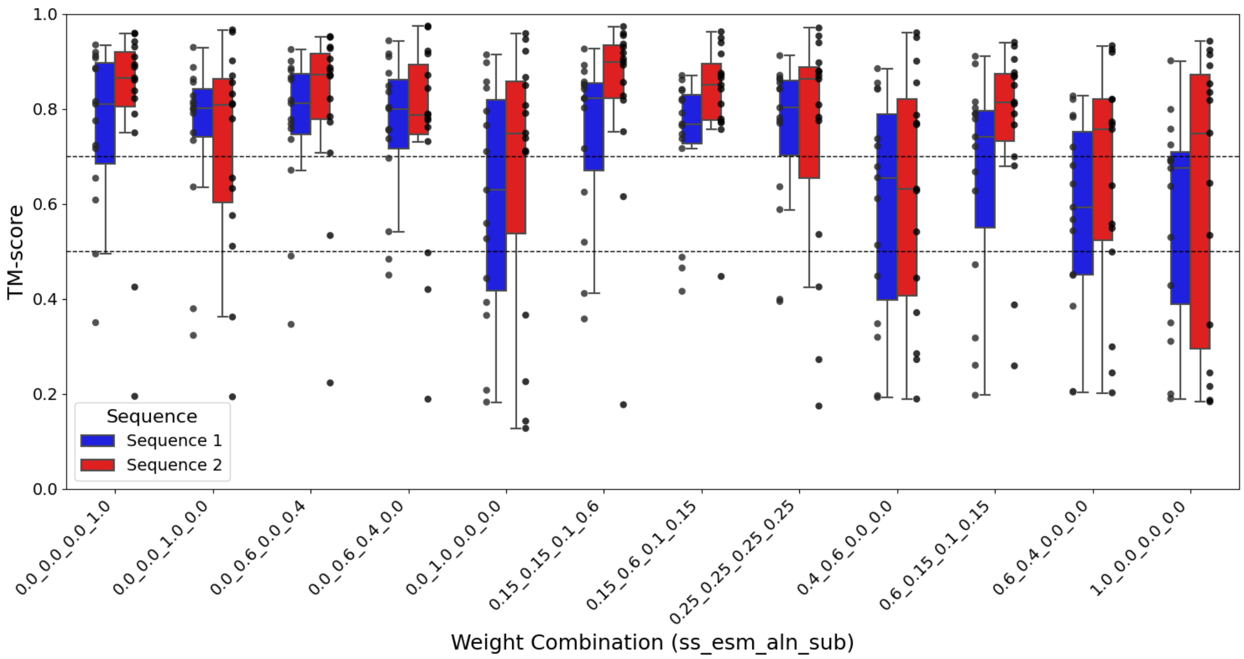


**Fig. S5. Convergent overlap optimization metrics across parameter weightings using 15 randomly selected protein pairs from SwissProt.**

Boxplots show the distribution of TM-scores (1) for each weight combination, with individual data points overlaid in black. TM-scores reflect the original sequence compared to the optimized convergent overlap sequence (both obtained using predictions via ColabFold (2)) Blue corresponds to Sequence 1 and red to Sequence 2. Dashed horizontal lines indicate TM-score thresholds at 0.5 (indicative of likely shared fold/topology) and 0.7 (indicative of a higher-confidence, near-native structural match) (3). Weight combinations are shown along the x-axis, with parameters ordered as ss (secondary structure), esm (ESM-2 contact map), aln (alignment), and sub (substitution). Only sequences whose terminal region exhibited a mean pLDDT ≥ 90 in the original AlphaFold2 prediction were included. This cutoff was applied to restrict the analysis to terminal regions predicted with very high confidence, thereby reducing potential noise from low-confidence residues.
